# Transposon mediated horizontal transfer of the host-specific virulence protein ToxA between three fungal wheat pathogens

**DOI:** 10.1101/671446

**Authors:** Megan C. McDonald, Adam P. Taranto, Erin Hill, Benjamin Schwessinger, Zhaohui Liu, Steven Simpfendorfer, Andrew Milgate, Peter S. Solomon

## Abstract

Most known examples of horizontal gene transfer (HGT) between eukaryotes are ancient. These events are identified primarily using phylogenetic methods on coding regions alone. Only rarely are there examples of HGT where non-coding DNA is also reported. The gene encoding the wheat virulence protein ToxA and surrounding 14 kb is one of these rare examples. *ToxA* has been horizontally transferred between three fungal wheat pathogens (*Parastagonospora nodorum, Pyrenophora tritici-repentis* and *Bipolaris sorokiniana*) as part of a conserved ∼14kb element, which contains coding and non-coding regions. Here we use long-read sequencing to define the extent of HGT between these three fungal species. Construction of near-chromosomal level assemblies enabled identification of terminal inverted repeats on either end of the 14kb region, typical of a Type II DNA transposon. This is the first description of *ToxA* with complete transposon features, which we call ToxhAT. In all three species, ToxhAT resides in a large (140-250 kb) transposon-rich genomic island which is absent in *toxA-* isolates. We demonstrate that the horizontal transfer of ToxhAT between *Pyrenophora tritici-repentis* and *P. nodorum* occurred as part of a large ∼80kb HGT which is now undergoing extensive decay. In contrast, in *B. sorokiniana* ToxhAT and its resident genomic island are mobile within the genome. Together these data provide insight into the non-coding regions that facilitate HGT between eukaryotes and the genomic processes which mask the extent of HGT between these species.

**IMPORTANCE:** This work dissects the tripartite horizontal transfer *of ToxA*; a gene that has a direct negative impact on global wheat yields. Defining the extent of horizontally transferred DNA is important because it can provide clues as to the mechanisms that facilitate HGT. Our analysis of *ToxA* and its surrounding 14kb suggests that this gene was horizontally transferred in two independent events, with one event likely facilitated by a Type II DNA transposon. These horizontal transfer events are now in various processes of decay in each species due to the repeated insertion of new transposons and subsequent rounds of targeted mutation by a fungal genome defense mechanism known as repeat induced point-mutation. This work highlights the role that HGT plays in the evolution of host adaptation in eukaryotic pathogens. It also increases the growing body of evidence that transposons facilitate adaptive HGT events between fungi present in similar environments and hosts.

**DATA AVAILABILITY:** All raw sequencing data is available under NCBI BioProject PRJNA505097.

The *P. nodorum* SN15 Whole Genome Shotgun project has been deposited at DDBJ/ENA/GenBank under the accession SSHU00000000. The version SSHU01000000 is described in this paper. The *P. nodorum* SN79-1087 Whole Genome Shotgun project has been deposited under the accessions CP039668-CP039689. The Whole Genome shotgun project and accession numbers for *B. sorokiniana* isolates are as follows: CS10; SRZH00000000, CS27; SRZG00000000, WAI2406; SRZF00000000, WAI2411; SRZE00000000. Transposon annotations, CS10 and CS27 gene annotations are available at https://github.com/megancamilla/Transposon-Mediated-transfer-of-ToxA

## INTRODUCTION

Horizontal gene transfer (HGT) is a mechanism whereby DNA from unrelated organisms is transferred in a non-Mendelian fashion (1). In proteobacteria HGT is thought to have occurred in over 75% of all protein families, making HGT one of the most important tools to facilitate adaptation to new, stressful environments (2, 3). This propensity to share DNA between species has been attributed to many human health issues, such as the rapid rise and spread of antibiotic resistance in hospitals (4). In eukaryotes, HGT was once thought to be a rare event and therefore not an important contributor to environmental adaptation. However, numerous studies have now shown that HGT between eukaryotes plays a very important role in adaptation, especially in the case of microbes that colonize a common host (5–12).

Among eukaryotic microbes, fungi are often used for kingdom-wide studies of adaptation, due to their relatively small genome size, importance in human and plant disease, and applications in food and biotechnology (5–7). Domesticated fungi, particularly those used in food production, are now being used as model organisms to understand the genetic basis of adaptation (8–10). On an evolutionary time scale these organisms have been subjected to a short but intense period of selection, which has dramatic effects on their preferred carbon and nitrogen sources, secondary metabolite production and many other physiological traits (10, 11). One emerging theme from these studies is that organisms which are common contaminants of the food making process are often donors of the genes that provide fitness advantages in these specialized environments. The reported HGT events are large and involve tens of thousands of bases of DNA, which remain over 90% identical between very distantly related species (8, 9). These HGTs contain both coding and non-coding regions which are stably integrated into the core nuclear genomes of the recipient species (8, 9). While the original reports suggested that these regions were important for adaptation to the domestic environment, the fitness advantage conferred by these genes had to be demonstrated in follow-up studies with knock-out strains (11, 12).

Rapid adaptation via HGT is not restricted to domesticated species, but there exist very few described instances where the horizontally transferred DNA is integrated into the core nuclear genome and remains highly identical outside of coding regions. One standout example is the virulence gene *ToxA* and the surrounding 11-12 kb, which to-date has been reported in three fungal wheat pathogens; *Parastagonospora nodorum, Pyrenophora tritici-repentis* and *Bipolaris sorokiniana* (13–16). While all three species belong to the same fungal order, the Pleosporales, they are distant relatives with several million years separating their speciation (13, 14). Similar to the domesticated fungi discussed above, this HGT event is hypothesized to be extremely recent as the average pairwise nucleotide identity across this 12 kb region remains greater than 92% (15). The *ToxA* gene itself remains identical between *P. tritici-repentis* and *B. sorokiniana* and only three nucleotides different from between *B. sorokiniana and P. nodorum* (15). The fitness advantage that *ToxA* confers has been demonstrated experimentally, whereby the presence of *ToxA* in a fungal isolate leads to faster development of necrotic lesions on wheat leaves (15, 16). This virulence function is genotype specific, as *ToxA* only causes necrosis on wheat lines that carry the susceptibility gene called *Tsn1* (19–21). In the absence of *Tsn1,* all three fungal species can still infect wheat due to the presence of other virulence genes (15, 17, 18).

Though *ToxA* confers a strong fitness advantage, this HGT event is not a fixed insertion and persists in all three pathogen populations as a presence/absence polymorphism (15, 19–21). The size of this presence/absence polymorphism has yet to be fully characterized. The presence of *ToxA* in different field populations around the world also varies dramatically, ranging from 6% to 97% presence in different pathogen field populations (20). The selective forces that increase the frequency of *ToxA* in some fungal populations and decrease it in others remain unknown. Studies that examined whether there was a positive correlation between the frequency of *ToxA* in fields planted with *Tsn1* (susceptible) wheat cultivars were inconclusive (22). For ease of reading we will use the notation *ToxA+* for isolates that contain the gene and *toxa-* for isolates that do not carry the gene.

Despite detailed knowledge on the molecular function of *ToxA* and its prevalence in fungal pathogen populations throughout the world, we still do not know the origins of this important virulence gene, nor the mechanisms that facilitated its transfer and stable integration into the genomes of these three pathogen species. In all three species there is clear evidence that *ToxA* is embedded in an AT-rich, repeat-dense region of the genome. AT-richness in these portions of the genome is driven by a fungal specific genome defense process known as Repeat Induced Point-Mutation (RIP) (23). RIP targets repeated sequences as small as 220 bp, mutating C:G to T:A, which introduces early stop codons in repeated DNA sequences (23–25). This mechanism is hypothesized to have evolved in some phyla of fungi to stop the spread of transposons or other self-copying elements within their genomes (24).

*ToxA* and its highly conserved flanking DNA provides a unique opportunity to dissect the integration of horizontally transferred DNA into the nuclear genomes of three fungal pathogens. To define the location and extent of each HGT event, we used long-read DNA sequencing to generate near-complete genome assemblies for several representatives from two of the three species, in addition to several other published assemblies (26, 27). We performed extensive *de novo* annotation of the repeat families in all three fungal species and manually annotated the region surrounding the *ToxA* gene. These assemblies and repeat-annotations resolve the genomic context in which the virulence gene is located and provide insights into potential mechanisms of HGT as well as the history of horizontal transfer events.

## RESULTS

### Long-read sequencing reveals a conserved Type II DNA transposon

The genomic location of *ToxA* is best described in *P. tritici-repentis,* where two long-read assemblies place this gene in the middle of Chromosome 06 (supercontig1.4) (21),(28). Several long-read assemblies have also been generated for *ToxA+ P. nodorum* isolates SN4 and SN2000, where *ToxA* is found on Chromosome 08 in both isolates (26). In addition to these publicly available assemblies we sequenced the *ToxA+* isolates *P. nodorum* SN15 and *B. sorokiniana* CS10 (original isolate name BRIP10943) with seven PacBio SMRT cells each resulting in approximately 500 thousand reads with an average read size of 10.6kb and 9.4kb, respectively. In addition to the two SMRT assemblies, we re-sequenced an additional four isolates with the Oxford Nanopore MinION. This included two *toxa-* isolates, *P. nodorum* isolate SN79-1087 and *B. sorokiniana* isolate CS27 (original isolate name BRIP27492a), as well as two additional *ToxA+ B. sorokiniana* isolates, WAI2406 and WAI2411. All isolates were *de novo* assembled using long-read data only. Short-read Illumina data was used to ‘polish’ the Nanopore *de novo* assemblies of CS27 and SN79-1087. A complete list of all isolates used in this study, their assembly method and assembly quality indicators are described in Table 1. Genome assembly accession numbers and additional information about the isolates are given in Table S1. *B. sorokiniana* chromosomes were ordered and named from largest to smallest based on the PacBio assembly of isolate CS10. Our *P. nodorum* contigs were named based on synteny alignments to the recently published assemblies from Richards *et al.* (26).

**TABLE 1:** Summary of genome assembly statistics for each isolate assembled in this study

| *Species* | *B. sorokiniana* |  |  |  | *P. nodorum* |  |
| --- | --- | --- | --- | --- | --- | --- |
| Isolate | CS10 | CS27 | WAl2406 | WAl2411 | SN15 | SN79-1087 |
| Raw Data | PacBio | Nanopore + Illumina | Nanopore | Nanopore | PacBio | Nanopore + Illumina |
| ToxA Genotype | *ToxA+* | *toxa-* | *ToxA+* | *ToxA+* | *ToxA+* | *toxa-* |
| Assembly Size (Mb)^ | 36.9 | 35.2 | 36.9 | 36.2 | 37.3 | 34.7 |
| # Contigs* | 22 | 19 | 21 | 21 | 26 | 23 |
| # Contigs w/both telomeres | 14 | NA | NA | NA | 18 | 13 |
| # Contig with 1 telomere | 3 | NA | NA | NA | 6 | 8 |
| BUSCO score<sup>++</sup> | 98.9 | 98.9 | 72.4 | 69.8 | 99.1 | 92.4 |

^ Size in Millions of base pairs
\* Nuclear contigs only
<sup>++</sup>Percentage of 1313 ascomycete single-copy orthologs found in assembly

Assembly quality was assessed using the Benchmarking Universal Single-Copy Orthologs (BUSCO) tool, which identifies fragmented, duplicated and missing genes from *de novo* assemblies (29, 30). The scores reported in Table 1 are the percentage of complete genes found in a set of 1313 BUSCO genes from the Ascomycota odb9 dataset. The number of complete genes is used as a proxy to estimate total genome completeness (29). The assembly completeness scores were much lower for Nanopore-only assemblies in isolates WAI2406 and WAI2411, where no short-read data was available for genome correction. For isolates CS27 and SN79-1087, available short-read data allowed correction of the assemblies, so that the number of complete BUSCO genes found exceeded 90%. Both PacBio assemblies, without short-read data, generated BUSCO scores greater than 98% (Table 1). For both PacBio assemblies, CS10 and SN15 the 6-bp telomeric repeat (TAACCC) was found on the ends of several contigs, summarized in Table 1. We could not identify telomeric repeats in the Nanopore assemblies for *B. sorokiniana* isolates. However, for the assembly of *P. nodorum* isolate SN79-1087 we were able to identify many contigs with telomeric repeats (Table 1).

We have previously reported the presence of a >92% identical 12kb region shared between all three species (15). This aligns with the original publication from Friesen *et al.* 2006, whereby a conserved 11kb element was reported between *P. tritici-repentis* and *P. nodorum*. In both *P. nodorum* SN15 and *B. sorokiniana* CS10 the chromosome that contains *ToxA* were assembled completely, with telomeric repeats on both ends. A self-alignment of this region in *B. sorokiniana* CS10 revealed intact, terminal inverted-repeats (TIRs) separated by 14.3 kb (Figure 1A). TIRs are structural features of Type II DNA transposons of the order “TIR”, which are required for excision by transposases (31). These TIRs were not identified in previous studies, which we explore further below (15, 18). The aligned TIRs are 74 bp and ∼92% identical (Figure 1B). We will hereafter refer to *ToxA* and the accompanying non-coding and coding DNA enclosed within these TIRs as “ToxhAT*”*. This name reflects the historical association of *ToxA* with the neighboring Type II hAT-like transposase gene (18).

**Figure 1:**
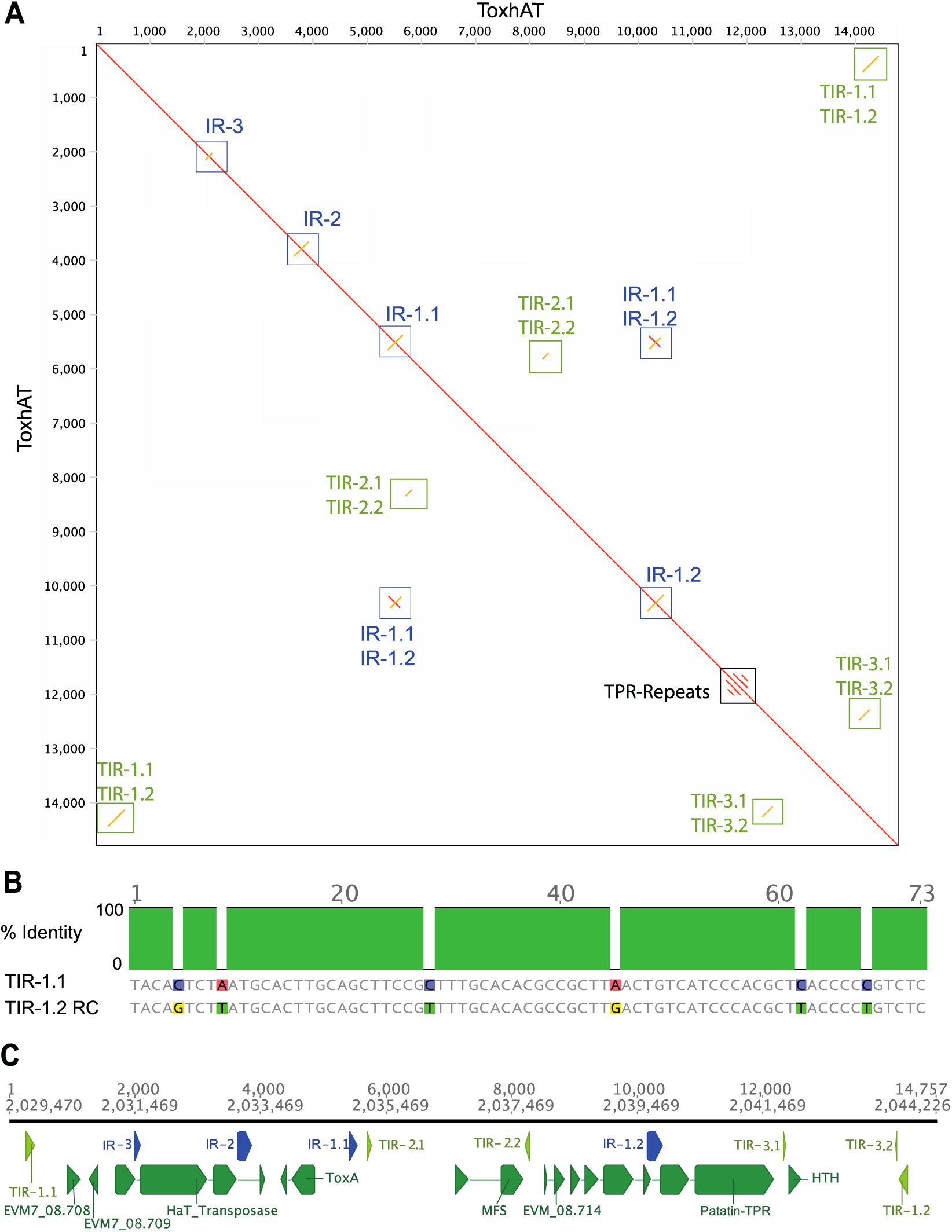
Characterization of ToxhAT in *B. sorokiniana* isolate CS10. A) Self alignment of ToxhAT drawn as a dot-plot. The red-line down the center shows a 1-to-1 alignment, yellow lines show inverse alignments. Terminal inverted repeats (TIRs) and inverted repeats (IRs) are boxed. The TPR-repeats are short tandem repeats found in the gene with the Patatin domain B) Alignment of the 74 bp TIR1.1 and the Reverse Complement (RC) of TIR1.2. Grey color indicates aligned positions that are identical between the two sequences. C) Manual annotation of coding regions within ToxhAT showing each open reading frame (green), inverted repeat (blue) and TIR (light green).

We annotated the coding regions within ToxhAT in *B. sorokiniana* isolate CS10 with both long and short read RNA-sequencing data. This annotation plus the self-aligned sequence revealed eight genes, three inverted repeats (IRs) and two additional internal TIRs (Figure 1C). Three annotated genes, *CS10_08.708*, *CS10_08.709* and *CS10_08.714* contain no known protein domains. Excluding *ToxA,* the remaining four genes had conserved domains, as identified by NCBI’s conserved domain database (Figure 1C). One gene contained a major facilitator superfamily (MFS) domain (accession:cl28910), which in yeast was shown to be a proton-coupled transporter of di- and tri-peptides (32). The largest CDS within ToxhAT contained two known protein domains, with a Patatin domain at the N-terminus (Accession:cd07216) followed by tetratricopeptide repeats (TPR, pfam13424) at the C-terminus. In fungi, proteins that contain these domains are recognized as members of the NOD-like receptor (NLR) family (33). Only a limited number of these proteins have been functionally studied in fungi, but they are broadly considered to be involved in self-recognition and immunity (33, 34). The fourth gene was flanked by its own set of TIRs and contained a helix-turn-helix (HTH) DNA binding domain (Accession:cl04999). This structure indicated that this CDS is likely a nested Type II transposase within ToxhAT (Figure 1C). This indicates that ToxhAT is a composite of at least two DNA transposons. Fragments of the eight open reading frames are also found in the other two species, *P. nodorum* and *P. tritici-repentis* (Fig. S1), however in these two species the 3’ end of ToxhAT is invaded by unique sequences (Fig. S1).

Using *B. sorokiniana* as a guide we were able to identify remnants of the ToxhAT TIRs in both *P. tritici-repentis* and *P. nodorum.* In *P. tritici-repentis* the 5’ TIR remains largely intact, whereas the 3’ TIR is enriched in C to T and G to A transitions characteristic of RIP (Fig. S2). For *P. nodorum* both the 5’ and 3’ TIR are enriched in RIP mutations, which without prior knowledge of the TIR location in *B. sorokiniana,* would be impossible to identify. In *P. tritici-repentis* and *P. nodorum* there were additional unique sequence insertions inside of ToxhAT (Figure 2, Fig. S1). Manual annotation of this unique sequence showed *P. tritici-repentis* 1C-BFP has a 3’ insertion of a Type II DNA transposon, a Tc1-mariner-like sequence, which interrupts ToxhAT separating the 5’-3’ TIRs by ∼20.2kb. In *P. nodorum* SN15 this element is interrupted by a different transposon, which resembles a Type I Long-terminal-repeat (LTR)-retrotransposon. The exact identity of this transposon was difficult to determine due to extensive RIP-like mutations in this sequence. This insertion separates the ToxhAT TIRS in *P. nodorum* by ∼25.6kb (Fig. S1). Despite these additional insertions the TIRs identified in *B. sorokiniana* are present in the other two species, though heavily mutated, indicating a common evolutionary origin for ToxhAT.

**Figure 2:**
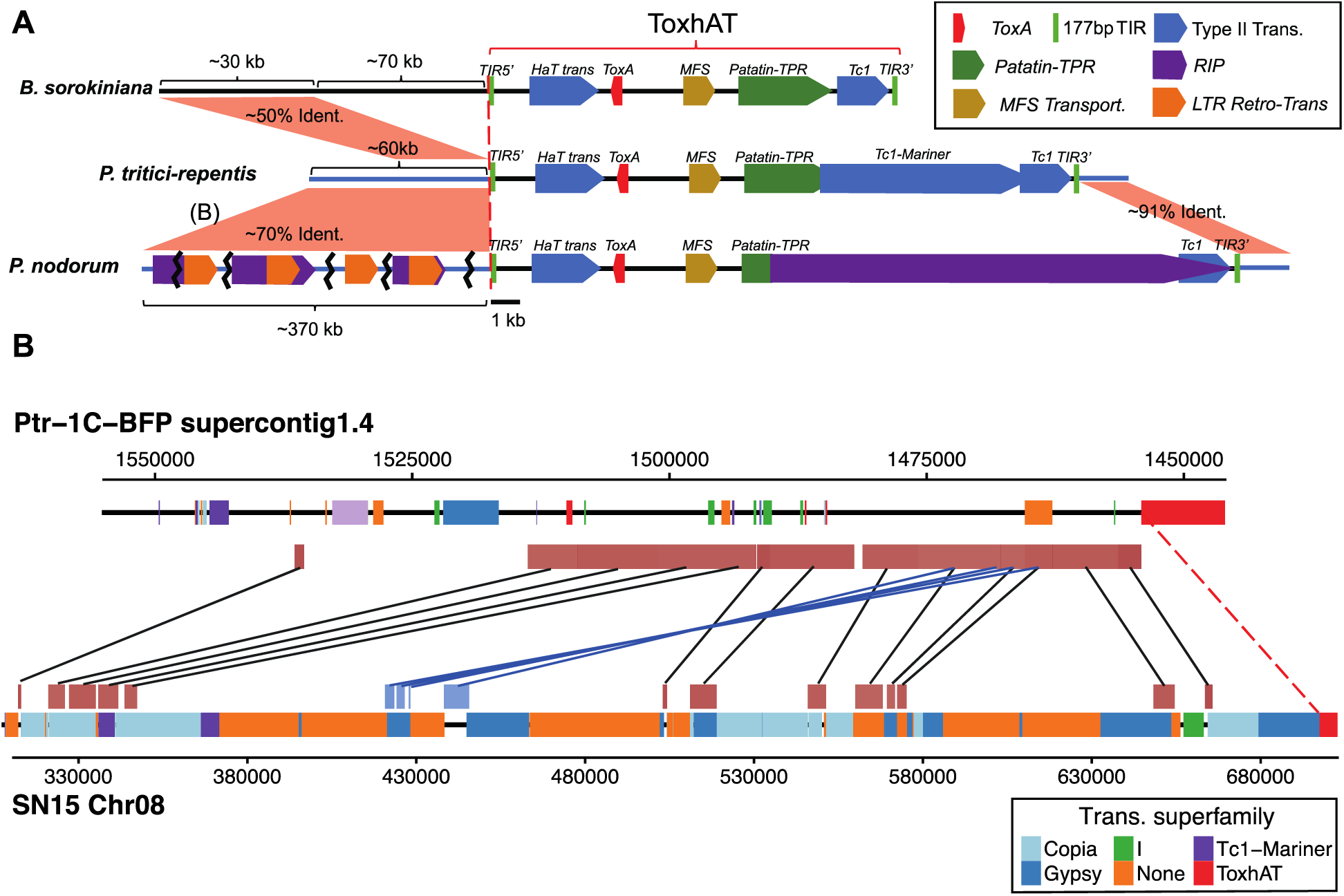
A) Overview of the ToxhAT in all three pathogen species. All features drawn to the right of the red-dashed line are drawn to scale as indicated by the black scale bar below. Features to the left of the red dashed line are not drawn to scale, with the relative size indicated with brackets. The opaque red rectangles drawn between the chromosomes outside of the TIRs show regions synteny as indicated by whole chromosome alignment. The approximate percent nucleotide identity is indicated within the red shading. Bracketed (B) in part A indicates the region shared between *P. nodorum* and *P. tritici-repentis,* which is drawn in part B) Close-up of whole chromosome alignment between *P. nodorum* and *P. tritici-repentis.* Chromosomes are drawn as thick black lines with positions of annotated transposons shown in colored blocks. Transposons are classified into superfamiles as indicated by the legend. The additional opaque red/blue boxes appearing above or below the chromosomes are nucleotide regions >70% identical identified by whole chromosome alignment with LASTZ. Black lines connect syntentic blocks aligned in the same direction, while blue lines connect inverted syntenic blocks.

### ToxhAT was transferred in two independent HGT events

Whole chromosomal alignments (WCA) between the ToxhAT containing chromosomes of *P. nodorum* and *P. tritici-repentis* revealed DNA with >70% sequence identity beyond the boundaries set by ToxhAT TIRs (Figure 2A, Fig S2). Pairwise alignments of these regions revealed that almost all polymorphisms were RIP-like (Fig. S3). Excluding the RIP-like mutations, the sequence identity nears 100% between the two species. Ten genes annotated in *Ptr 1C-BFP,* PTRG-04890-04909, are all present in *P. nodorum* SN15 upstream of the 5’ ToxhAT TIR. However, in *P. nodorum*, each of these is a pseudogene due to RIP and therefore have not been annotated in any assembly (Fig. S3) (19, 26). Furthermore, five of these 10 genes, PTRG-04891-04895, are duplicated and found in inverse orientation within the *P. nodorum* SN15 assembly (Figure 2B, blue boxes). In *P. tritici-repentis* these 10 genes are on a contiguous piece of DNA that extends 61.2 kb upstream of the ToxhAT 5’-TIR. The total length of near identical sequence shared between these two species is ∼80kb, which includes 61.2 kb upstream and 1.7 kb downstream of ToxhAT. In *P. nodorum* SN15 the 61.2 kb shared with *P. tritici-repentis* is present but highly fragmented across Chromosome 08, spanning nearly ∼370kb (Figure 2B). This data demonstrates that a specific HGT event occurred between *P. tritici-repentis* and *P. nodorum* that included ToxhAT and a large segment of surrounding DNA.

WCA between *P. tritici-repentis* and *B. sorokiniana* also revealed ∼30kb outside of ToxhAT that was ∼50% identical and partially overlapped the DNA shared *between P. tritici-*repentis *and P. nodorum* (Figure 2A, inclusive of the genes PTRG-04892-04900). This indicated that a region outside of ToxhAT was potentially also horizontally transferred between *B. sorokiniana* and *P. tritici-repentis.* However, a pairwise alignment of this region showed no evidence of extensive RIP-like mutations that could account for the pairwise nucleotide differences observed between these two species (Fig. S4). Furthermore, we could identify the same region in the *toxA-* isolate *B. sorokiniana* ND90Pr as well as in other closely related *Bipolaris spp.* (Fig. S5). We conclude that this region was not part of a horizontal transfer of ToxhAT into *B. sorokiniana* and is instead a region of synteny between the two species.

### Individual components of ToxhAT are found in other Pleosporales

After careful manual annotation of ToxhAT, we conducted a full *de novo* repeat prediction and annotation with the REPET pipeline (35, 36). The total proportion of each genome annotated as repeats is shown in Fig. S6. The non-redundant repeat library generated by REPET included the manually annotated ToxhAT transposon from *B. sorokiniana* CS10, named DTX-comp_CS10_RS_00, and a second near full-length version from *P. nodorum*, named DTX-incomp-chim_SN2000-L-B14-Map1. These two sequences were used to identify all instances of ToxhAT within each genome listed in Table 1 and *P. tritici-repentis* 1C-BFP. 195 instances of ToxhAT were annotated in these seven isolates, 183 (∼94%) of which we were able to successfully align to the CS10 ToxhAT sequence (Fig. S7). This alignment showed distinct areas within ToxhAT that were found in high abundance within these genomes, overlapping primarily with *CS10_08.708*, *CS10_08.709*, and the Patatin-like gene (Fig. S7). A summary of the total number of identified ToxhAT instances is summarized in Table 2. These data show that most annotations are fragments with the median length ranging from 176 to 295 bp, approximately one percent of the total length of ToxhAT.

**Table 2:** Summary of REPET identified ToxhATs in each isolate.

| Species | Isolate | N | Max Length<br>ToxhAT (bp) | Min Length<br>ToxhAT (bp) | Average<br>Length (bp) | Median<br>Length (bp) |
| --- | --- | --- | --- | --- | --- | --- |
| *B. sorokiniana* | CS10 | 42 | 14079 | 28 | 831 | 251 |
|  | CS27 | 22 | 1552 | 26 | 437 | 179 |
|  | WAI2406 | 35 | 14082 | 28 | 935 | 225 |
|  | WAI2411 | 41 | 14072 | 28 | 891 | 250 |
| *P. tritici-repentis* | Ptr_1CBPF | 29 | 20213 | 32 | 1081 | 188 |
| *P. nodorum* | SN15 | 18 | 11153 | 54 | 1219 | 295 |
|  | SN79 | 8 | 537 | 99 | 266 | 261 |

The large number of partial ToxhAT annotations in *toxa-* isolates suggested some regions may be repetitive elements independent of ToxhAT. To investigate this further we performed tBLASTn queries on the NCBInr database and the Dothideomycetes genomes available at JGI MycoCosm. In both searches the hAT transposase, MFS transporter and Patatin-TPR genes had over 500 partial hits with an e-value less than 1e-10 (Figs. S8A and S8B). Within the JGI Dothideomycetes database search, *CS10_08.708* had 139 hits, *CS10_08.709* had 85 hits, and the *Tc1* transposase had 263 hits (e-value <1e-10). The small gene *CS10_08.714* had the fewest number of hits with only four in total across both databases (excluding known instances of ToxhAT). Hits for the *ToxA* gene itself were mostly distant homologs (<50% identical), previously described as *ToxA-like* or *ToxA\** in various *Bipolaris spp.* (37). A short summary of the top hits from both database searches are presented in Table 3.

**Table 3:**
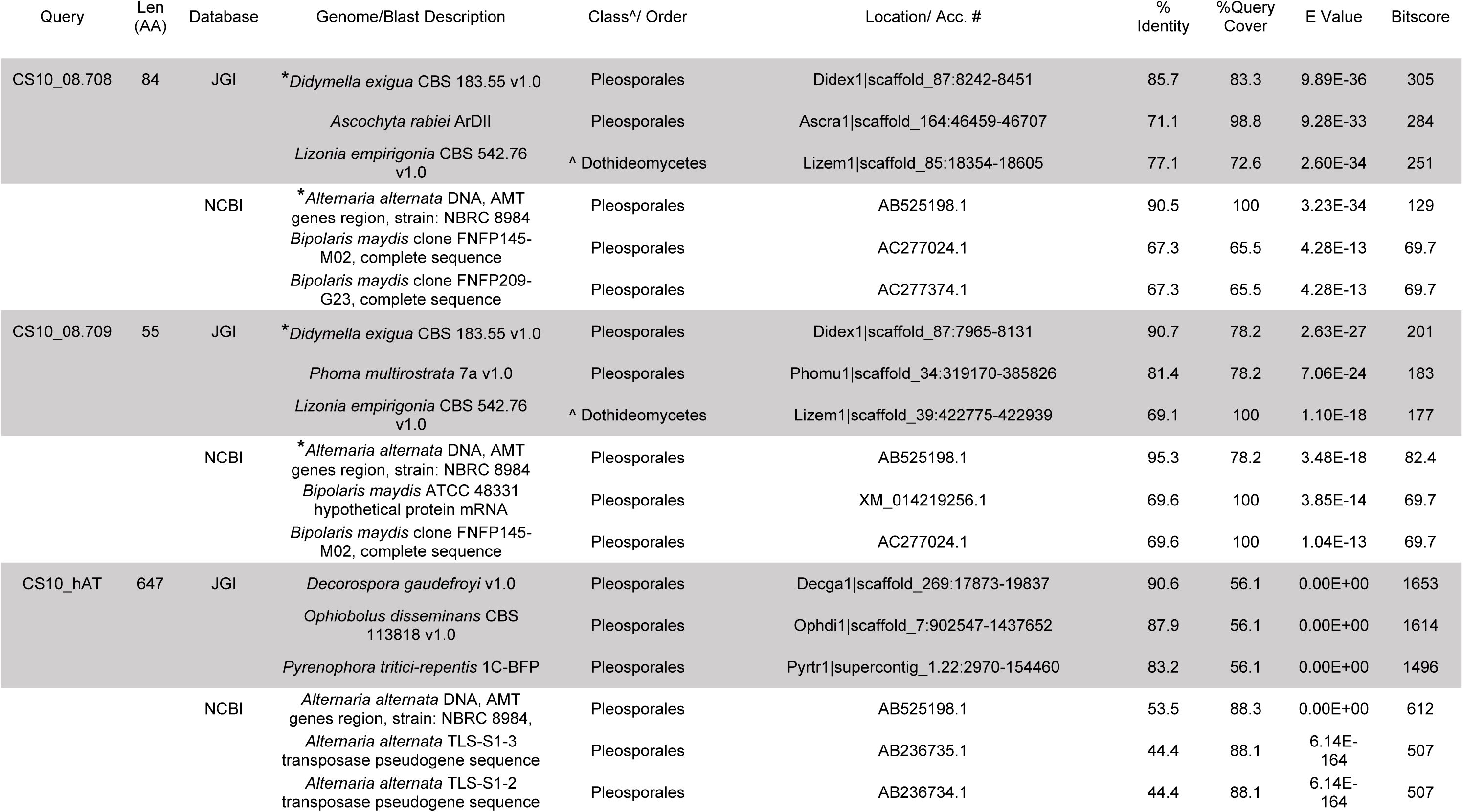
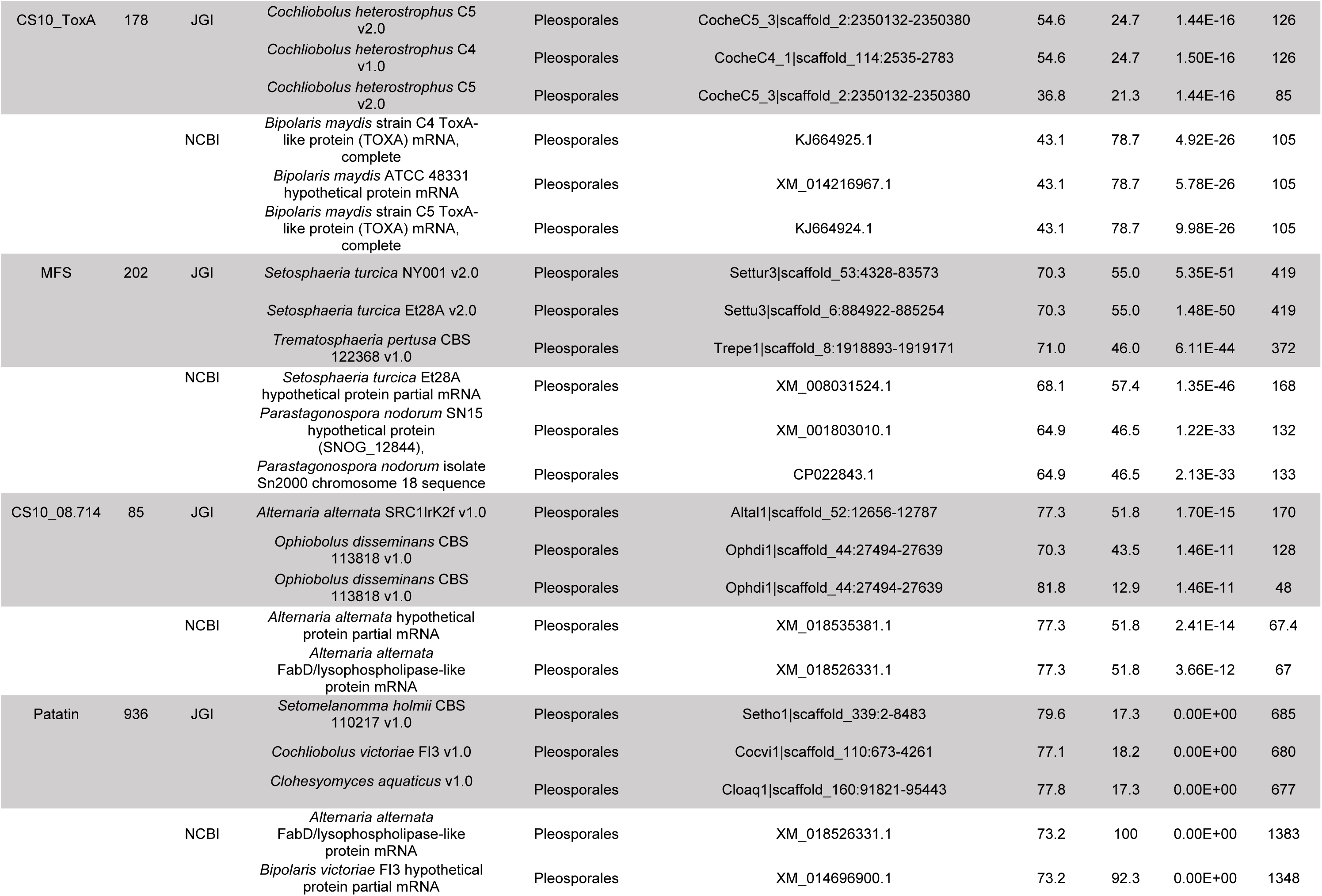
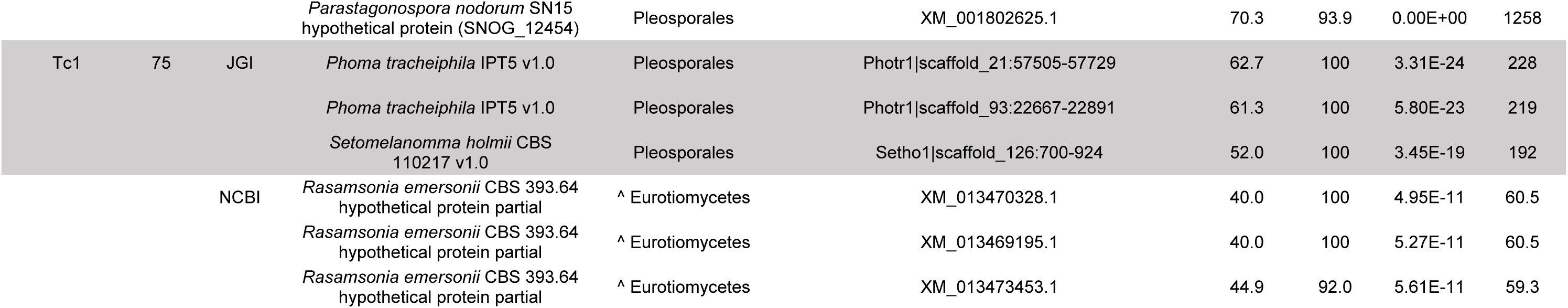
Summary of top BLAST hits excluding known examples of ToxhAT in *P. tritici-repentis, P. nodorum* and *B. sorokiniana.* Only the Top 3 BLAST hits are shown sorted according to bitscore, which is a measure of pairwise sequence similarity. ^*^Indicates hits of interest with >85% pairwise nucleotide identity and >78% query coverage. ^^^ Class is indicated for species outside of the Dothideomycetes or when the order is unknown

| Query | Len<br>(AA) | Database | Genome/Blast Description | Class^/ Order | Location/ Acc. # | %<br>Identity | %Query<br>Cover | E Value | Bitscore |
| --- | --- | --- | --- | --- | --- | --- | --- | --- | --- |
| CS10_08.708 | 84 | JGI | * *Didymella exigua* CBS 183.55 v1.0 | Pleosporales | Didex1\|scaffold_87:8242-8451 | 85.7 | 83.3 | 9.89E-36 | 305 |
|  |  |  | *Ascochyta rabiei* ArDII | Pleosporales | Ascra1\|scaffold_164:46459-46707 | 71.1 | 98.8 | 9.28E-33 | 284 |
|  |  |  | *Lizonia empirigonia* CBS 542.76 v1.0 | ^ Dothideomycetes | Lizem1\|scaffold_85:18354-18605 | 77.1 | 72.6 | 2.60E-34 | 251 |
|  |  | NCBI | * *Alternaria alternata* DNA, AMT genes region, strain: NBRC 8984 | Pleosporales | AB525198.1 | 90.5 | 100 | 3.23E-34 | 129 |
|  |  |  | *Bipolaris maydis* clone FNFP145-M02, complete sequence | Pleosporales | AC277024.1 | 67.3 | 65.5 | 4.28E-13 | 69.7 |
|  |  |  | *Bipolaris maydis* clone FNFP209-G23, complete sequence | Pleosporales | AC277374.1 | 67.3 | 65.5 | 4.28E-13 | 69.7 |
| CS10_08.709 | 55 | JGI | * *Didymella exigua* CBS 183.55 v1.0 | Pleosporales | Didex1\|scaffold_87:7965-8131 | 90.7 | 78.2 | 2.63E-27 | 201 |
|  |  |  | *Phoma multirostrata* 7a v1.0 | Pleosporales | Phomu1\|scaffold_34:319170-385826 | 81.4 | 78.2 | 7.06E-24 | 183 |
|  |  |  | *Lizonia empirigonia* CBS 542.76 v1.0 | ^ Dothideomycetes | Lizem1\|scaffold_39:422775-422939 | 69.1 | 100 | 1.10E-18 | 177 |
|  |  | NCBI | * *Alternaria alternata* DNA, AMT genes region, strain: NBRC 8984 | Pleosporales | AB525198.1 | 95.3 | 78.2 | 3.48E-18 | 82.4 |
|  |  |  | *Bipolaris maydis* ATCC 48331 hypothetical protein mRNA | Pleosporales | XM_014219256.1 | 69.6 | 100 | 3.85E-14 | 69.7 |
|  |  |  | *Bipolaris maydis* clone FNFP145-M02, complete sequence | Pleosporales | AC277024.1 | 69.6 | 100 | 1.04E-13 | 69.7 |
| CS10_hAT | 647 | JGI | *Decorospora gaudefroyi* v1.0 | Pleosporales | Decga1\|scaffold_269:17873-19837 | 90.6 | 56.1 | 0.00E+00 | 1653 |
|  |  |  | *Ophiobolus disseminans* CBS 113818 v1.0 | Pleosporales | Ophdi1\|scaffold_7:902547-1437652 | 87.9 | 56.1 | 0.00E+00 | 1614 |
|  |  |  | *Pyrenophora tritici-repentis* 1C-BFP | Pleosporales | Pyrtr1\|supercontig_1.22:2970-154460 | 83.2 | 56.1 | 0.00E+00 | 1496 |
|  |  | NCBI | *Alternaria alternata* DNA, AMT genes region, strain: NBRC 8984, | Pleosporales | AB525198.1 | 53.5 | 88.3 | 0.00E+00 | 612 |
|  |  |  | *Alternaria alternata* TLS-S1-3 transposase pseudogene sequence | Pleosporales | AB236735.1 | 44.4 | 88.1 | 6.14E-164 | 507 |
|  |  |  | *Alternaria alternata* TLS-S1-2 transposase pseudogene sequence | Pleosporales | AB236734.1 | 44.4 | 88.1 | 6.14E-164 | 507 |
| CS10_ToxA | 178 | JGI | *Cochliobolus heterostrophus* C5 v2.0 | Pleosporales | CocheC5_3\|scaffold_2:2350132-2350380 | 54.6 | 24.7 | 1.44E-16 | 126 |
|  |  |  | *Cochliobolus heterostrophus* C4 v1.0 | Pleosporales | CocheC4_1\|scaffold_114:2535-2783 | 54.6 | 24.7 | 1.50E-16 | 126 |
|  |  |  | *Cochliobolus heterostrophus* C5 v2.0 | Pleosporales | CocheC5_3\|scaffold_2:2350132-2350380 | 36.8 | 21.3 | 1.44E-16 | 85 |
|  |  | NCBI | *Bipolaris maydis* strain C4 ToxA-like protein (TOXA) mRNA, complete | Pleosporales | KJ664925.1 | 43.1 | 78.7 | 4.92E-26 | 105 |
|  |  |  | *Bipolaris maydis* ATCC 48331 hypothetical protein mRNA | Pleosporales | XM_014216967.1 | 43.1 | 78.7 | 5.78E-26 | 105 |
|  |  |  | *Bipolaris maydis* strain C5 ToxA-like protein (TOXA) mRNA, complete | Pleosporales | KJ664924.1 | 43.1 | 78.7 | 9.98E-26 | 105 |
| MFS | 202 | JGI | *Setosphaeria turcica* NY001 v2.0 | Pleosporales | Settur3\|scaffold_53:4328-83573 | 70.3 | 55.0 | 5.35E-51 | 419 |
|  |  |  | *Setosphaeria turcica* Et28A v2.0 | Pleosporales | Settu3\|scaffold_6:884922-885254 | 70.3 | 55.0 | 1.48E-50 | 419 |
|  |  |  | *Trematosphaeria pertusa* CBS 122368 v1.0 | Pleosporales | Trepe1\|scaffold_8:1918893-1919171 | 71.0 | 46.0 | 6.11E-44 | 372 |
|  |  | NCBI | *Setosphaeria turcica* Et28A hypothetical protein partial mRNA | Pleosporales | XM_008031524.1 | 68.1 | 57.4 | 1.35E-46 | 168 |
|  |  |  | *Parastagonospora nodorum* SN15 hypothetical protein (SNOG_12844), | Pleosporales | XM_001803010.1 | 64.9 | 46.5 | 1.22E-33 | 132 |
|  |  |  | *Parastagonospora nodorum* isolate Sn2000 chromosome 18 sequence | Pleosporales | CP022843.1 | 64.9 | 46.5 | 2.13E-33 | 133 |
| CS10_08.714 | 85 | JGI | *Alternaria alternata* SRC1lrK2f v1.0 | Pleosporales | Altal1\|scaffold_52:12656-12787 | 77.3 | 51.8 | 1.70E-15 | 170 |
|  |  |  | *Ophiobolus disseminans* CBS 113818 v1.0 | Pleosporales | Ophdi1\|scaffold_44:27494-27639 | 70.3 | 43.5 | 1.46E-11 | 128 |
|  |  |  | *Ophiobolus disseminans* CBS 113818 v1.0 | Pleosporales | Ophdi1\|scaffold_44:27494-27639 | 81.8 | 12.9 | 1.46E-11 | 48 |
|  |  | NCBI | *Alternaria alternata* hypothetical protein partial mRNA | Pleosporales | XM_018535381.1 | 77.3 | 51.8 | 2.41E-14 | 67.4 |
|  |  |  | *Alternaria alternata* FabD/lysophospholipase-like protein mRNA | Pleosporales | XM_018526331.1 | 77.3 | 51.8 | 3.66E-12 | 67 |
| Patatin | 936 | JGI | *Setomelanomma holmii* CBS 110217 v1.0 | Pleosporales | Setho1\|scaffold_339:2-8483 | 79.6 | 17.3 | 0.00E+00 | 685 |
|  |  |  | *Cochliobolus victoriae* FI3 v1.0 | Pleosporales | Cocvi1\|scaffold_110:673-4261 | 77.1 | 18.2 | 0.00E+00 | 680 |
|  |  |  | *Clohesyomyces aquaticus* v1.0 | Pleosporales | Cloaq1\|scaffold_160:91821-95443 | 77.8 | 17.3 | 0.00E+00 | 677 |
|  |  | NCBI | *Alternaria alternata* FabD/lysophospholipase-like protein mRNA | Pleosporales | XM_018526331.1 | 73.2 | 100 | 0.00E+00 | 1383 |
|  |  |  | *Bipolaris victoriae* FI3 hypothetical protein partial mRNA | Pleosporales | XM_014696900.1 | 73.2 | 92.3 | 0.00E+00 | 1348 |
|  |  |  | *Parastagonospora nodorum* SN15<br>hypothetical protein (SNOG_12454) | Pleosporales | XM_001802625.1 | 70.3 | 93.9 | 0.00E+00 | 1258 |
| Tc1 | 75 | JGI | *Phoma tracheiphila* IPT5 v1.0 | Pleosporales | Photr1\|scaffold_21:57505-57729 | 62.7 | 100 | 3.31E-24 | 228 |
|  |  |  | *Phoma tracheiphila* IPT5 v1.0 | Pleosporales | Photr1\|scaffold_93:22667-22891 | 61.3 | 100 | 5.80E-23 | 219 |
|  |  |  | *Setomelanomma holmii* CBS<br>110217 v1.0 | Pleosporales | Setho1\|scaffold_126:700-924 | 52.0 | 100 | 3.45E-19 | 192 |
|  |  | NCBI | *Rasamsonia emersonii* CBS 393.64<br>hypothetical protein partial | ^ Eurotiomycetes | XM_013470328.1 | 40.0 | 100 | 4.95E-11 | 60.5 |
|  |  |  | *Rasamsonia emersonii* CBS 393.64<br>hypothetical protein partial | ^ Eurotiomycetes | XM_013469195.1 | 40.0 | 100 | 5.27E-11 | 60.5 |
|  |  |  | *Rasamsonia emersonii* CBS 393.64<br>hypothetical protein partial | ^ Eurotiomycetes | XM_013473453.1 | 44.9 | 92.0 | 5.61E-11 | 59.3 |

The highest-identity hits were in a single *Alternaria alternata* strain: NBRC 8984 (AB525198.1), for two contiguous open reading frames, *CS10_08.708*, *CS10_08.709*. The pairwise identity for these two genes exceeded 90% across and are co-localized in *A. alternata* NBRC 8984. This strain was also the highest-identity hit in NCBI for the hAT-transposase. *A. alternata* is a well-known plant pathogen, that has a broad host range and is also a member of the Pleosporales. In the JGI database a fungal isolate collected from leaf litter, *Didymella exigua* CBS 183.55 v1.0, also had co-localized hits for *CS10_08.708*, *CS10_08.709* with identity greater than 85%, indicating that these two predicted genes may be a single repetitive unit. This species is also classified in the Pleosporales (38). The third species identified with >90% sequence identity for the hAT transposase was *Decospora gaudefroyi,* again classified in the Pleosporales as a salt-tolerant marine fungus (39). Overall, the large number and breadth of hits across different fungal species confirmed our hypothesis that the individual coding regions of ToxhAT are part of repetitive elements found broadly within the Pleosporales. This indicates a common evolutionary origin of these repetitive coding regions within this fungal Order.

### The presence/absence polymorphism of *ToxA* is much larger than the extent of HGT

Above we showed the extent of shared DNA varies between different pairwise comparisons of the three species harboring ToxhAT. This does not address the unknown size of the presence/absence polymorphism maintained in these species. To investigate this, the homologous *toxa-* and *ToxA+* chromosomes were aligned (Figure 3). For *P. tritici-repentis,* where no long-read data for a *toxa-* isolate was available, short reads from isolate DW5 were aligned against the assembly of PTR1C-BFP. The large spike in coverage for DW5 within the deleted region corresponds to a 7kb TIR transposon most likely from the Tc1-Mariner superfamily, which is found near ToxhAT (DTX-incomp-chim_Ptr-L-B62-Map1_reversed). The chromosomes containing *ToxA* in the isolates *P. nodorum* SN15 and *B. sorokiniana* CS10 both contain telomeric repeats and are complete chromosomes. This shows that in all three species the absence polymorphism spans several thousand kb (Figure 3). Using the last known homologous regions from the whole chromosome alignments, the absence alleles in *B. sorokiniana* CS27*, P. nodorum* SN79-1087 and *P. tritici-repentis* DW5 were estimated to span ∼239 kb, ∼467 kb and ∼150 kb, respectively.

**FIGURE 3:**
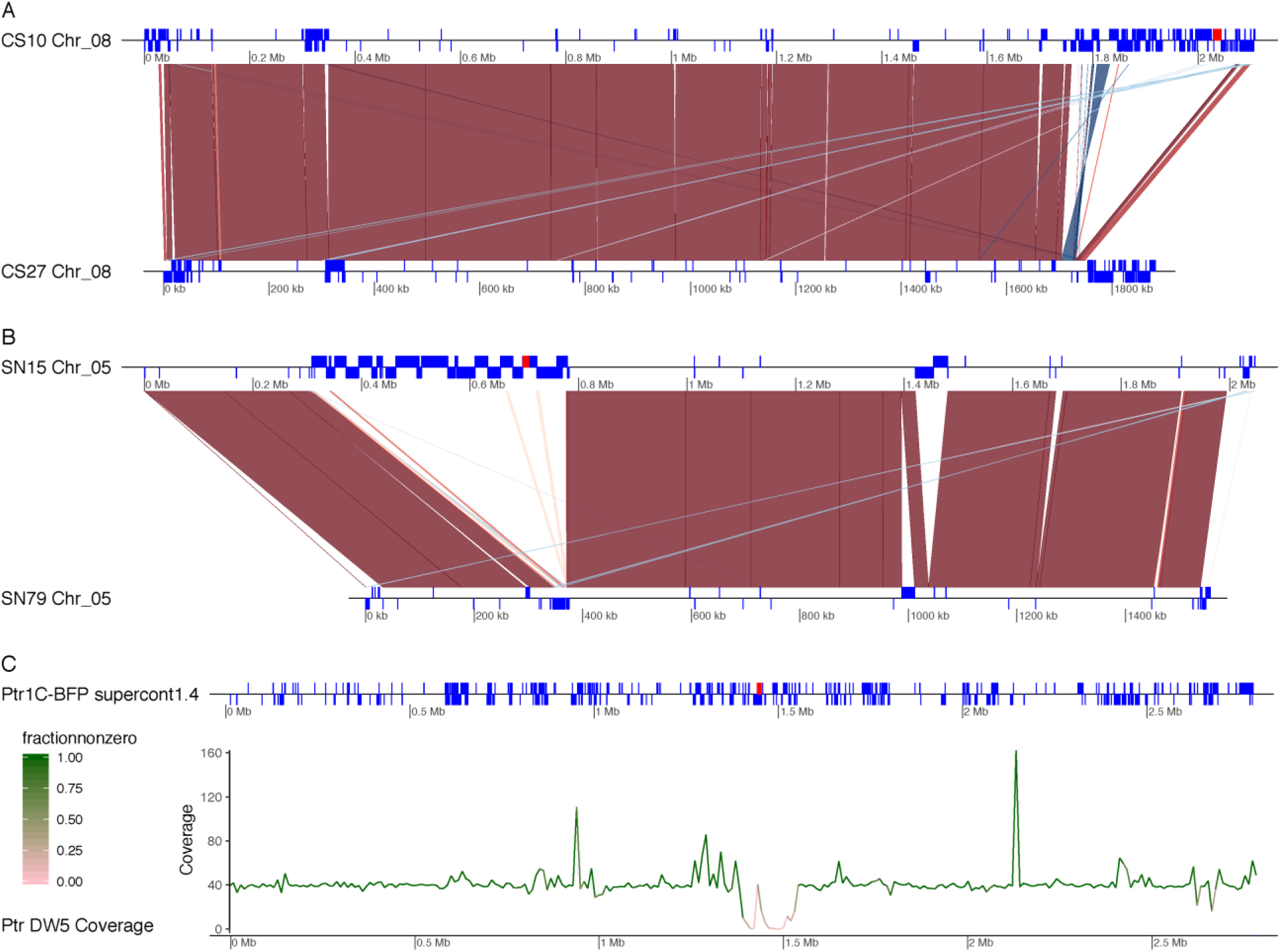
Genomic context of the ToxhAT containing region (red box) in each of the three species in comparison to *toxa-* isolates. A) Lastz alignment of the homologous scaffold between *B. sorokiniana ToxA+* isolate CS10 and *toxa-* isolate CS27. Blue blocks drawn on the scaffold maps (black lines) represent the location of annotated transposons within each genome. Red ribbons drawn between the two isolates represent syntenic alignments found in Lastz greater than 70% identity and 2kb in length. Blue ribbons drawn between the two isolates show inversions between the two genomes. B) Lastz alignment of the homologous scaffold between *P. nodorum ToxA+* isolate SN15 and *toxa-* isolate SN79-1087. Same coloring scheme as in part A. C) Isolate *Ptr*1C-BFP with repeat regions shown in the blue blocks along the scaffold map (black line). Below is the average Illumina coverage for 10 kb windows across the chromosome. The color of the line corresponds to the proportion of bases within the 10kb window that have non-zero coverage.

To compare the size of the presence/absence polymorphism of *ToxA* with two other well characterized necrotrophic effectors we examined the location of *SnTox3* and *SnTox1* in *P. nodorum* SN15 (40). These two effectors also exist as a presence/absence polymorphism in this species but have no known history of HGT. In isolate SN15, Sn*Tox3* is the last annotated gene on Chromosome 11 and the absence polymorphism is approximately the 7kb tail of this chromosome. This absence encompasses the annotated SN15 genes, SNOG_08981 (*Tox3*) -SNOG-08984 (Fig. S9A). The end of Chromosome 11 in the SN79-1087 (which lacks *SnTox3*) assembly contains telomeric repeats (data not shown), which suggests that the missing 7kb is not due to an incompletely assembled chromosome. The absence allele of *SnTox1* is even smaller, spanning ∼3kb on *SN15’s* Chromosome 6. At this locus there is a unique insertion of ∼1.3 kb which is only present in SN79-1087 (Fig. S9B). This data demonstrates that the absence allele of the horizontally transferred *ToxA* is much larger than absence alleles of other known effectors and highlights potential genome instability after HGT events.

### Evidence of mobility of ToxhAT in *Bipolaris sorokiniana*

The intact TIRs found in *B. sorokiniana* CS10 suggested that ToxhAT in this species may remain mobile. To investigate this, we re-sequenced two additional *ToxA+* isolates of *B. sorokiniana* (WAI2406 and WAI2411). In both genomes ToxhAT was found in different genomic locations when compared to isolate CS10, where ToxhAT is located near the end of Chromosome 08 (Figure 4A). For WAI2406, ToxhAT and the surrounding ∼200kb of repeat-rich DNA was found imbedded in the middle of Chromosome 01 (Figure 4A-C). This has led to an increase in size of WAI2406’s Chromosome 01 which is ∼4.0 Mbp compared to CS10’s PacBio assembled Chromosome 01 of ∼3.8 Mbp. To confirm that this translocation was not a mis-assembly, we aligned the corrected Nanopore reads from Canu to both the CS10 and WAI2406 assemblies (Figure 4B-C). These reads aligned with a slope of 1, to Chromosome 01 in isolate WAI2406 with single reads clearly spanning the break-points on both sides of the translocation. These same reads from isolate WAI2406 did not align well to Chromosome 08 in isolate CS10.

**FIGURE 4:**
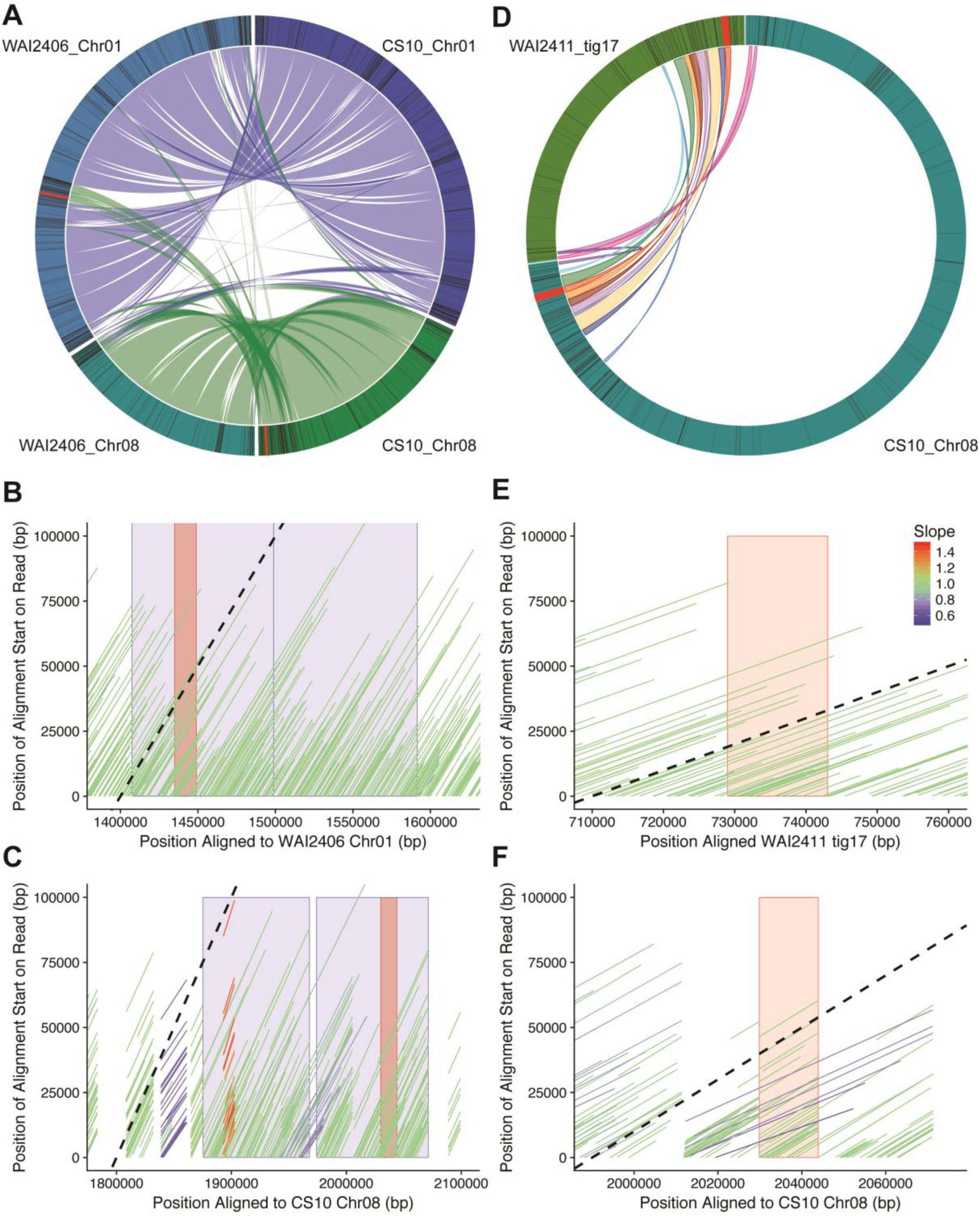
ToxhAT within *B. sorokiniana* is mobile in two distinct ways. A) Shows the alignment of CS10’s Chromosomes 01 and 08 against WAI2406 homologous chromosomes. The ToxhAT (red line/box drawn on outer circle) is located on Chr01 in WAI2406 along with a large region of repeat-rich DNA (black boxes in outer circle). B) The corrected and trimmed WAI2406 Nanopore reads aligned to the *de novo* of itself. The black dotted line shows a slope=1, which is indicative that the read aligned base-per-base against the chromosome shown on the x-axis. Reads with a slope different from 1 are reads that have been mapped dis-continuously (i.e. with large insertions or deletions). The blue blocks show the translocated DNA, found in Chromosome 08 in CS10 but in Chromosome 01 in WAI2406. The red box indicates the position of ToxhAT. C) The same reads as in B but aligned to Chromosome 08 of CS10. Note read alignment is not continuous and breaks at the translocation edges. D) Shows the alignment of Chromosome 08 from CS10 against tig17 from WAI2411. E) Same as shown in B but with isolate WAI2411, red block shows the position of ToxhAT F) Same as shown in C but with isolate WAI2411, note no reads with a slope of 1 extend beyond the ToxhAT itself (red box).

For isolate WAI2411, ToxhAT assembled to a small contig (∼776kb) that has homology to the PacBio assembled CS10’s Chromosome 02 and Chromosome 08 (data not shown). While it remains unclear if this small scaffold is a part of Chromosome 02 or Chromosome 08, the flanking DNA on either side of ToxhAT was conserved but shuffled in order (Figure 4D). ToxhAT was found to be inverted and in a different position in comparison to Chromosome 08 in CS10. The breakpoints of the inversion were precisely from TIR to TIR of ToxhAT. Again, we aligned the corrected nanopore reads to isolate WAI2411 (self) and isolate CS10. The self-alignment showed reads that clearly crossed the break-points of the inversion/transposition (Figure 4E), however these same reads did not map contiguously to the CS10 assembly (Figure 4F). As a secondary confirmation we used BLASTn to identify all reads that contain *ToxA* and generated a multiple sequence alignment (Fig. S10). Twenty-nine single-molecule reads were aligned that spanned ToxhAT and continued into both flanking regions. These flanking regions aligned well to WAI2411’s contig 17 confirming the inversion of ToxhAT precisely at the TIRs (Figure 4E). We postulated that this inversion could be an active transposition event, in which case there may be signature target-site duplications (TSDs) made by the transposase. However, this inversion nor any other instance of ToxhAT in any sequenced isolate was flanked by identifiable target site duplications.

## DISCUSSION

As shown in previous studies the *ToxA* gene and surrounding non-coding DNA is highly conserved between the three species (15, 18, 27). We extend this knowledge by defining flanking TIRs that give this region structural features of a Type II DNA transposon, defined herein as ToxhAT. These TIRs and the enclosed 14.3 kb are conserved in all three fungal species, indicating ToxhAT has a common evolutionary origin prior to HGT. Homologous DNA shared between *P. nodorum* and *P. tritici-repentis* outside of these TIRs indicates that the HGT event between these two species was not ToxhAT alone but included ∼63 kb of flanking DNA. In contrast homology between these two species and *B. sorokiniana* breaks precisely at the ToxhAT TIRs. Within *B. sorokiniana,* there is strong evidence that ToxhAT and the surrounding repeat-rich DNA remains mobile within the genome. The individual coding regions found within ToxhAT appear to be part of repetitive elements in other Dothideomycetes. However, the breadth of hits, gives no indication of any single species that could have assembled ToxhAT as a unit before horizontal transfer between these three wheat pathogens.

### ToxhAT is structured like a Type II DNA transposon which remains mobile in *B. sorokiniana*

By identifying conserved TIRs in all three pathogens we describe a unit of DNA that has the structural features of a Type II DNA TIR transposon. In *B. sorokiniana* isolate WAI2411, ToxhAT itself is inverted precisely at the TIRs. This inversion bounded precisely by these structural features suggested that this putative transposon may remain active. However, extensive searches of the flanking DNA in all three species did not reveal target site duplications (TSDs) typical of other TIR transposases (41, 42). TSDs are created by a transposase when it cuts at its target site, usually a short sequence ranging from two to eleven bases (31). Most transposases make an un-even cut, which after DNA repair, leads to a duplication of the target site on either side of the inserted transposon (31). The absence of TSDs in WAI2411 suggests that the inversion seen in isolate WAI2411 was not facilitated by an active transposition event. An alternative mechanism that could explain the inversion is intra-chromosomal recombination between these structural features (23). In the absence of TSDs we consider this mechanism a strong alternative to explain the movement of ToxhAT in WAI2411.

While WAI2411 showed a relatively small genomic rearrangement, the other re-sequenced *B. sorokiniana* isolate WAI2406 contained a large segmental movement of 200 kb from one chromosome to another. Similar inter-chromosomal translocations were observed in the fungal plant pathogens *Verticillium dahliae, Magnaporthe oryzae* and *Colletotrichum higginsianum* (43–45). While all three studies demonstrate that translocations occur in regions where transposons are prevalent, only the study by Faino *et al.* (2016) in the asexual species *V. dahliae,* was able to show homologous transposons/DNA sequence at the translocation breakpoints. These breakpoints in *V. dahliae* implicate homologous recombination as the mechanism underpinning genome plasticity in this species (46). In *M. oryzae* and *C. higginsianum,* transposons are associated with translocation events but it remains unclear if these translocations occurred over regions with high sequence identity (43, 45). Similarly, in *B. sorokiniana* whole chromosome alignment of Chromosome 08 and Chromosome 01 in isolate WAI2406 did not show high sequence identity in the regions outside of the translocation breakpoints on either chromosome. However, the regions near these breakpoints are enriched in transposon annotations. A key difference between the three fungal species examined in this study and *Verticillium* is their sexual lifecycle. Meiotic recombination could obscure the precise translocation boundaries and is also a pre-requisite for RIP. We postulate that in sexual fungal species, AT-rich regions, with otherwise limited sequence identity, may undergo “homologous” inter-chromosomal recombination events like those observed in *B. sorokiniana*. Supporting this hypothesis is a recent study on *Epichloë festucue* that leveraged HiC data to build a contact map of DNA within the nucleus (47). The HiC sequencing technique shows the frequency at which different regions of a genome interact with each other in the nucleus (48). For example, DNA fragments that are physically close to each other, on the same chromosome, have a higher frequency of interaction when compared to sequences on different chromosomes (47). The study by Winter *et al.* (2018) showed that AT-rich regions, usually heavily RIPed transposon islands, on different chromosomes had significantly higher interactions with each other when compared to non-AT rich regions (47). These data suggest that in sexual fungi, where RIP is active, AT-rich islands are associated with each other in 3D space. We postulate that AT-rich regions in *B. sorokiniana,* like *E. festucue*, also associate with each other within the nucleus, which provides the physical proximity required for inter-chromosomal recombination events.

The opportunity to examine inter-chromosomal transfer of ToxhAT extends beyond *B. sorokiniana.* Chromosomal movement of the *ToxA* gene was also observed in *P. tritici-repentis* (49). In this study, pulse-field gel electrophoresis followed by southern hybridization was used to show that *ToxA* was found on chromosomes of different sizes in *P. tritici-repentis.* Going further, the authors showed that in at least one isolate *ToxA* was on a different chromosome when compared to the *ToxA* location in reference isolate 1C-BFP (49). While this study probed for the *ToxA* gene alone, we consider it likely that ToxhAT or a larger chromosomal segment was translocated, similar to what was observed in *B. sorokiniana.* Further long-read assembly coupled with HiC data from both *P. tritici-repentis* and *B. sorokiniana,* ideally from a sexual cross of two previously sequenced isolates, is required to systematically reconstruct the level of sequence identity or other genome features that facilitate inter-chromosomal translocations.

### ToxhAT resides in an accessory genomic region in all three species

The analysis of the syntenic relationship between *ToxA*+ and *toxA*-chromosomes within each species showed that the absence of ToxhAT is coincident with the absence of large, >100kb, chromosomal segments. The DNA composition of these regions fits well within the definition of “lineage specific” or “accessory” regions described in pathogenic fungi, where virulence genes are found nested within transposon-rich regions of the genome (50–53). This genome structure is hypothesized to facilitate the rapid adaptation of fungal pathogenicity genes, often referred to as the “two-speed” or “two-compartment” genome (54, 55). These data further show that the absence polymorphism does not coincide with the exact boundaries of HGT. This is most clearly seen in *P. nodorum* where fragments of the horizontally transferred DNA are scattered across a 300kb region. However, this entire region, extending well beyond the horizontally transferred (HT) fragments, is missing from the *toxA-* isolate SN79-1087. This leads to an interesting question about whether the horizontal acquisition of ToxhAT precipitated the expansion of transposons within this region. Our data for *SnTox3* and *SnTox1* in *P. nodorum* suggests that this may be the case, whereby the absence alleles span only a few kilobases. Unfortunately, similar comparisons in *B. sorokiniana* were not possible due to a lack of known effectors and in *P. tritici-repentis,* where the only other known effector, *ToxB,* is present in multiple copies in the genome (56). Intra-chromosomal recombination is also a possible mechanism to generate these large absence polymorphisms. In many model organisms, such as Drosophila, yeast and human cell lines, large segmental deletions were facilitated by ectopic recombination between tandemly arrayed repeat sequences (57, 58). This mechanism is particularly interesting in the context of HGT, as these recombination events often result in the formation of circular extra-chromosomal DNA (59, 60).

### The origins of ToxhAT and mechanism for HGT remain obscure

Since the discovery of *ToxA* in the genome of *P. nodorum,* the evolutionary origin of this gene has been a topic of debate (18, 21, 61, 62). To date, *P. nodorum* remains the species with the highest known *ToxA* sequence diversity. This diversity underpins the prevailing hypothesis that *ToxA* has had the most time to accumulate mutations and therefore has resided in the genome of this organism longest (18, 61). The discovery of *ToxA* in *B. sorokiniana* and characterization of the conserved 74 bp TIRs in all three species, strongly suggests that ToxhAT, has a single evolutionary origin in all three species. In *B. sorokiniana* the ToxhAT TIRs define the exact boundaries of the HGT event, where sequence identity with the other two species abruptly ends. In contrast, the HGT between *P. tritici-repentis* and *P. nodorum,* included ToxhAT and 63kb of flanking DNA. In *P. tritici-repentis* 1C-BFP, this flanking DNA remains contiguous, however in *P. nodorum* this same DNA is fragmented and partially duplicated across a 370kb island of RIPed transposable elements. Together these data suggest that ToxhAT was horizontally transferred in two separate events both with and without flanking DNA.

We propose two opposing models to explain the HGT of ToxhAT between the three species. In the first model we assume a population genetic perspective where the longer the DNA has resided in the genome the more fragmented and dispersed the HT DNA will become. In this model *P. nodorum* would be the donor of ToxhAT along with the flanking DNA to *P. tritici-repentis.* This is based on our observation that the flanking DNA outside of ToxhAT is highly fragmented and duplicated in *P. nodorum,* indicating a longer period of time to accumulate these changes. This model assumes that there remains a donor isolate, in *P. nodorum* which once had a contiguous stretch of DNA as observed in *P. tritici-repentis.* In this model we also postulate that ToxhAT is most recently acquired by *B. sorokiniana* due to its relatively compact form and conservation of structural features. Again, for this model to hold, we must assume that an intact form of ToxhAT exists in the population of *P. nodorum* or *P. tritici-repentis* that could act as a donor to *B. sorokiniana.* Our second model assumes that changes can be accumulated more rapidly in transposon rich regions, and therefore are not a good indication of evolutionary time. In this model the intact version of ToxhAT observed in *B. sorokiniana* would represent an ancestral version. This is the minimal unit of HT DNA, bounded by conserved TIRs. In this model we propose the first HGT event is from *B. sorokiniana* to *P. tritici-repentis,* based on the identical *ToxA* sequence that they share. Then in a second HGT event, *P. tritici-repentis* would be the donor of the large segment of DNA inclusive of *ToxhAT* to *P. nodorum*. In *P. nodorum* the HT DNA flanking ToxhAT was subject to rapid decay and duplication, due to its proximity to transposable elements. This model does not provide a good explanation of why the rapid decay is not also observed in the other two species. Population scale long-read sequencing of *ToxA+* isolates from all three species is required to comprehensively test the validity of either of these models.

While we have presented two models above which describe the history of HT between these species alone, the BLAST searches conducted on the coding regions annotated in *B. sorokiniana*, indicate that there are other species which may harbor highly identical components of ToxhAT. One standout isolate is the *A. alternate* strain NBRC8984, which carries two genes that are 90% and 95% identical to *CS10_08.708* and *CS10_08.709*, respectively. Similar to their arrangement in ToxhAT, these two coding sequences also neighbor each other in *A. alternata* NBRC8984. While neither of these predicted genes have any known functional domains, they are by far the most similar hit to ToxhAT in a species that is not reported to carry *ToxA*. These high identity hits also included some non-pathogenic and marine species also found within the Pleosporales. Collectively, the coding regions within ToxhAT had hundreds of hits across species representing several hundred million years of evolution. However none of these coding regions have been characterized as repetitive or classified in a transposon family. Despite the ancient evolutionary history of transposons, the vast majority of described DNA transposons with TIRs are classified into only ten superfamilies (31, 63). Our detailed characterization of ToxhAT highlights an opportunity to characterize novel transposon superfamilies in non-model fungi.

### Towards a mechanism; flanking non-coding DNA provides clues

The tBLASTn results coupled with a detailed structural characterization of ToxhAT, suggests that it is a mosaic of repetitive coding regions. We propose that ToxhAT was transferred horizontally as, or by, a transposon with the fitness advantage of *ToxA* fixing these HGT events in three wheat-infecting species. Similarly, the horizontally transferred regions in the cheese-making *Pencillium spp.* were flanked by unusual i non-LTR retrotransposons (11). Horizontal transfer of transposons (HTT) has been widely reported in eukaryotes since the early discovery of P elements in *Drosophila* (64, 65). The literature on this topic however seems to clearly divide HGT from HTT as two separate phenomenon, the latter being much more common (66–68). Recent reports of the HTT between insects has used non-coding regions flanking horizontally transferred genes to demonstrate that a viral parasite, with a broad insect host range, is the vector for the horizontally transferred DNA (69). This study highlights how insights from non-coding regions can bring these studies closer to a mechanistic understanding of the HGT event (69, 70). Other studies which report HGT, or HT secondary metabolite clusters into and between fungal species often rely on phylogenetic methods on coding regions alone to detect these events (71–74). While these studies focus on the biological significance of the coding regions, clues to a possible mechanism may remain in the surrounding non-coding DNA. One limitation from early genome assemblies, was the inability to correctly assemble highly-repetitive regions. Here we demonstrate with two long-read sequencing technologies, it is possible to assemble very large repetitive regions heavily affected by RIP. These assemblies allow us to look at the non-coding regions that may be important for the movement or integration of HT DNA. Further population scale long-read sequencing will enable further refinement of the role that transposons play in facilitating adaptive gene transfer.

## Materials and Methods

### Fungal culture and DNA extraction

Fungal cultures were grown on V8-PDA media at 22°C under a 12hr light/dark cycle (75). Cultures ranging in age from 5-10 days were scraped from the surface of agar plates using a sterile razor blade into water. These harvested cultures were freeze-dried for 48 hours to remove all water. High molecular weight (HMW) DNA was extracted using a C-TAB phenol/chloroform method modified from Fulton *et al.* (1995) (76). Full details of our protocol including gel images of final DNA are available at http://dx.doi.org/10.17504/protocols.io.hadb2a6. DNA size was assessed by pulse field electrophoresis and DNA purity by examining 260/280 and 230/280 UV absorbance ratios on the Nanodrop (Thermo Scientific, USA). The total quantity of DNA was measured using the Qubit fluorometer (Life Technologies, USA).

### Genome sequencing and Assembly

#### PacBio DNA sequencing

All raw data generated from this study are deposited on NCBI BioProject ID PRJNA505097. Individual isolate accession numbers are listed in Table S1. Isolates SN15 (*P. nodorum*) and CS10 (*B. sorokiniana,* BRIP10943) were sequenced using Pacific Biosciences SMRT sequencing. 15-20kb Genomic P6 SMRT cell library preps were made at the Ramaciotti Centre for Genomics (UNSW Sydney, Australia). Each library was sequenced on 7 SMRT cells using the P6-C4 chemistry on the Pacbio RSII instrument (Pacific Biosciences, USA). Libraries were size selected on a Blue Pippin from 15-50kb. Each isolate genome was assembled *de novo* using Canu v1.5 and a minimum read length of 2000 bp (77). The canu setting “genomeSize” was set to 41Mb for isolate SN15 and 35 Mb for isolate CS10. Canu assemblies were further corrected using the SMRT Analysis package v2.3.0. First the raw Pacbio reads were mapped to the *de novo* assembly with blaser with the following settings: --seed=1 -- minAccuracy=0.75 --minLength=500 --forQuiver --algorithmOptions=’ -minMatch 12 -bestn 1 - minPctIdentity 65.0 --hitPolicy=randombest. The resulting bam file was used as input for Quiver to call a new consensus sequence with the following settings: makeVcf=True, makeBed=True, enableMapQVFilter=T, minConfidence=40, minCoverage-10, diploidMode=False (78).

#### Nanopore DNA sequencing

Resequencing of *B. sorokiniana* isolates CS27 (BRIP27), WAI2406 and WAI2411 and *P. nodorum* isolate SN79-1087 was performed on Oxford Nanopore’s MinIon sequencer. R9.4 flow cells were used for sequencing and the 1D library kit SQK-LSK08 was used to prepare the libraries according to the manufacturer’s protocol. All DNA samples were purified using Agencourt AMPure beads prior to starting the 1D library preparation (Beckman coulter, Inc. CA, USA). Genomes were assembled with Canu v1.5 or v1.6 with a minimum read length of 5kb (77). *De novo* genome assemblies were corrected using the trimmed reads output from Canu. Trimmed reads were mapped to the genome with Minimap2 followed by correction with Racon (79). The output consensus sequence from Racon was used as input for additional corrections steps performed iteratively up to five times. SN79-1087 and CS27 assemblies were further refined using the software Pilon, this correction was also performed iteratively up to five times (80). lllumina data from for SN79-1087 was taken from Syme et al. (2013) and for BRIP27 from McDonald et al. (2017).

### RNA-sequencing to aid annotation of long-read assemblies

*Bipolaris sorokiniana* isolate CS10 was cultured for 10 days in a range of liquid growth media – V8-juice broth, potato dextrose broth (PDB) (75); PDB supplemented with the epigenetic modifier, 5-azacytidine (150µM) (81); minimal media (82); wheat extract-supplemented minimal media; Fries 3; and Fries 3-modified media (83). Mycelia was harvested and total RNA extracted with the ZymoResearch™ Fungal/Bacterial RNA Miniprep kit. RNA quality assessment (Agilent Bioanalyzer), library preparation (strand-specific TruSeq v3) and Illumina RNA-sequencing (MiSeq, 150 bp single-end reads) were performed at the Ramaciotti Centre for Genomics (UNSW Sydney, Australia).

Long-read RNA-seq data was also generated using the Nanopore MinION. Total RNA extracted from mycelia cultured in Fries 3 was enriched for mRNA with the NEBNext® Oligo d(T) magnetic beads and concentrated with Agencourt® RNAclean® XP magnetic beads (Beckman Coulter, Inc. CA, USA). Direct RNA-seq and cDNA-PCR libraries were generated with ONT SQK-RNA001 and SQK-PCS108 library prep kits, respectively, and sequenced with R9.4 flow cells. Reads were basecalled with Albacore v2.0.2, quality filtered with Nanofilt and error-corrected using the CS10 genome sequence with Proovread (default settings) (84). Error-corrected reads were filtered for reads corresponding to full-length transcripts using SQANTI, run with default settings (85).

### Annotation of long-read assemblies

Illumina RNA-seq data was used for gene prediction in both CS10 and CS27 *B sorokiniana* isolates. Reads were trimmed with Trimmomatic v0.32 (parameters: -phred 33, ILLUMINACLIP TruSeq3-SE.fa:2:30:10, SLIDINGWINDOW:4:20, LEADING:20, TRAILING:20 and MINLEN:75) and aligned to the genomes using STAR v2.5 (parameters: --alignIntronMin 10, --alignIntronMax 1000, -- twoPassMode Basic), before transcript assembly using StringTie v1.3.3 (default parameters, except for --f 0.3). StringTie transcripts were filtered for high-quality ORFs using TransDecoder v5.0.2 (86–88).

Transcripts and aligned Illumina RNA-seq reads from all culture conditions were pooled for gene prediction. Pooled transcripts were used for gene prediction with CodingQuarry v2.0 (self-training Pathogen mode; default parameters) (89). Aligned reads and protein sequences from *P. nodorum* were used as evidence for gene prediction by BRAKER v2.0 (nondefault parameters --fungus, --prg gth) and GeMoMa v4.33 (default parameters) (90, 91). Gene predictions were combined with a nonredundant (nr) set of reviewed fungal Uniprot protein sequences and high-quality StringTie transcripts using EVidenceModeler (EVM), which generated a weighted consensus of all predicted gene models (Haas et al., 2008). Evidence sources were weighted accordingly: CodingQuarry > BRAKER ≥ GeMoMa > StringTie transcripts > nr fungal proteins. An assembly of MinION RNA-seq reads concordant with EVM gene models and StringTie transcripts was generated using PASA; this was used to update the EVM models (e.g. correcting intron-exon boundaries) and annotate UTRs (92). The completeness of these gene models was assessed using the BUSCO ascomycete database (29, 30).

### Annotation of Transposons

Transposons were identified *de novo* using the TEdenovo pipeline distributed as part of the REPET package v2.5 (35, 36). Long-read assemblies from the following species and isolates were used for *de novo* discovery; *P. nodorum,* isolates SN2000 and SN4 assembled by Richards *et al.* (2018), SN15 (this study), *P. tritici-repentis* 1-C-BFP, *B. sorokiniana* isolate CS10. Repbase v20.05 was used as the reference transposon database. TEs from each genome were combined into a common database according to the parameters set in Tedenovo_six_dnLibTEs_90_92.cfg. After combining TEs we manually added the coordinates of ToxhAT and named this TE “DTX-comp_CS10_RS_00”. Finally, TE’s were annotated in each genome listed in Table 1 with the common TE database using the TEannot pipeline and setting available in TEannot.cfg. file available at https://github.com/megancamilla/Transposon-Mediated-transfer-of-ToxA. Transposons were automatically classified into three letter codes based on the Wicker nomenclature (31, 93). (The final REPET summary files including repeat annotations for each genome can be found online at: https://github.com/megancamilla/Transposon-Mediated-transfer-of-ToxA.

Inverted repeats and TIRs within flanking the ToxA gene and surrounding DNA were identified in Geneious with the Dotplot (Self) viewing tool, which is based on the EMBOSS 6.5.7 tool dotmatcher (http://emboss.sourceforge.net/apps/release/6.5/emboss/apps/dotmatcher.html) (94). The specific settings required to reproduce the line plot shown in Figure 1 are as follows: Reverse complement=yes, Score Matrix=exact, window size=100, Threshold=75 and Tile Size=1000.

### Whole chromosome alignments

Initial whole genome alignments (WGA) were conducted using Lastz v1.02.00 or Mauve as implemented in Geneious v.9.1.8 (96–98). To obtain a clean gff or tab delimited file of high identity segments we developed Mimeo, which parses the alignment output of LASTZ (95–97). Mimeo and a description of its full features can found here: https://github.com/Adamtaranto/mimeo. WGA of ToxA+ and toxA-isolates was performed with LASTZ as implemented in mimeo with the following settings: mimeo-map --minIdt 60 --minLen 100 --maxtandem 40 –writeTRF. Candidate transfer regions between the three species were also identified by WGA preformed with LASTZ, as above with mimeo-map -–minIdt 70 --minLen 100. All alignments were inspected manually in Geneious v9.1.8. Chromosomal alignments were plotted in R v3.5.2 using the package genoPlotR (98). The corrected and trimmed nanopore reads output by Canu were used to align back to the de novo assemblies. These reads were mapped to the assembly with Minimap2 with the following settings: minimap2 -x map-ont -a. The output pairwise alignment file (paf) was modified in R for plotting. The complete R_markdown document which includes all code to reproduce Figures 2, 3B and 4 can be found at https://github.com/megancamilla/Transposon-Mediated-transfer-of-ToxA.

### BLAST searches

All open reading frames identified within ToxhAT in isolate CS10 were translated using Geneious v9.18. These amino acid sequences were used as queries in tBLASTn searches on NCBInr database and JGI MycoCosm with the following settings: Blosum62 Matrix, Gap Costs: Existence 11 extension 1, e-value max 1e-10, max hits=500 (Last Accessed 10^th^ Nov., 2018) (https://genome.jgi.doe.gov/programs/fungi/index.jsf) (99).

## Supplemental Material

**Table S1**: Genome assembly accession numbers and additional information about the isolates.: https://github.com/megancamilla/Transposon-Mediated-transfer-of-ToxA/tree/master/S1_GenomeStats

File S2: Gene annotations for CS10 and CS27: https://github.com/megancamilla/Transposon-Mediated-transfer-of-ToxA/tree/master/S2_Bipolaris_Gff3

File S3: REPET Annotation files: https://github.com/megancamilla/Transposon-Mediated-transfer-of-ToxA/S3_REPET_Files

File S4: Rmkd for Figure construction: https://github.com/megancamilla/Transposon-Mediated-transfer-of-ToxA/S4_Fig_Rmkds

## Acknowledgements

MCM would like to acknowledge The Sun Foundation’s Peer Prize for Women in Science for support to sequence additional *ToxA* isolates. EH acknowledges The Grains and Research Development Corporation: project UHS11002. MCM, AM, SS and PSS also acknowledge The Grains and Research Development Corporation for the collection of isolates: projects DAN00203 and DAN00177.

